# D-cysteine is an endogenous regulator of neural progenitor cell dynamics in the mammalian brain

**DOI:** 10.1101/2021.07.05.451211

**Authors:** Evan R. Semenza, Maged M. Harraz, Efrat Abramson, Adarsha P. Malla, Chirag Vasavda, Moataz M. Gadalla, Michael D. Kornberg, Solomon H. Snyder, Robin Roychaudhuri

## Abstract

D-amino acids are increasingly recognized as important signaling molecules in the mammalian central nervous system. However, the D-stereoisomer of the amino acid with the fastest *in vitro* spontaneous racemization rate, cysteine, has not been examined in mammals. Using chiral high-performance liquid chromatography and an stereospecific luciferase assay, we identify endogenous D-cysteine in the mammalian brain. We identify serine racemase (SR), which generates the NMDA glutamate receptor co-agonist D-serine, as a candidate biosynthetic enzyme for D-cysteine. Levels of D-cysteine are enriched over twentyfold in the embryonic mouse brain compared to the adult. D-cysteine reduces the proliferation of cultured mouse embryonic neural progenitor cells (NPCs) by approximately 50%, effects not shared with D-serine or L-cysteine. The antiproliferative effect of D-cysteine is mediated by the transcription factors FoxO1 and FoxO3a. The selective influence of D-cysteine on NPC proliferation is reflected in overgrowth and aberrant lamination of the cerebral cortex in neonatal SR knockout mice. Finally, we perform an unbiased screen for D-cysteine-binding proteins in NPCs by immunoprecipitation with a D-cysteine-specific antibody followed by mass spectrometry. This approach identifies myristoylated alanine-rich C-kinase substrate (MARCKS) as a putative D-cysteine-binding protein. Together, these results establish endogenous mammalian D-cysteine and implicate it as a physiologic regulator of NPC homeostasis in the developing brain.

## INTRODUCTION

Biochemical stereospecificity permits the extraordinary precision of enzymatic reactions and receptor-ligand interactions. On account of the selective incorporation of L-amino acids into proteins, D-amino acids were long considered “unnatural” isomers, occurring endogenously only in microorganisms and invertebrates (**1**). However, since the first report of free endogenous D-aspartate in human tissues in 1986 (**2**), D-amino acids have come to be appreciated as physiologic signaling molecules in mammals, particularly in the central nervous system (**3**).

The physiological roles of D-aspartate and D-serine, the first two D-amino acids established in mammals (**2, 4**), remain the most well-characterized. Both molecules act at N-methyl-D-aspartate-type glutamate receptors (NMDARs), with D-serine serving as the major endogenous ligand for the co-agonist (“glycine”) site (**5**), while D-aspartate binds the glutamate site in addition to neuroendocrine functions (**6–8**). Metabolism of diverse D-amino acids is mediated by analogous pathways, as both D-serine and D-aspartate are generated by pyridoxal 5’-phosphate-dependent enzymes, serine racemase (SR) and aspartate racemase, respectively (**9, 10**), and degraded by the flavin adenine dinucleotide-dependent oxidoreductases D-amino acid oxidase (DAAO) and D-aspartate oxidase (**11–13**).

The D-stereoisomers of a majority of proteogenic amino acids have been identified in the mammalian brain (**14**). Despite this widespread research into the presence and function of mammalian D-amino acids, the endogenous D-stereoisomer of the amino acid with the fastest *in vitro* spontaneous racemization rate, cysteine (**15**), has not been examined. In the current study we identify endogenous D-cysteine in the mammalian brain. D-cysteine is greatly enriched in embryonic mouse brain, with levels exceeding 4 mM, and is physiologically regulated by SR. D-cysteine, but not D-serine, reduces proliferation of cultured embryonic neural progenitor cells through Akt-mediated disinhibition of the transcription factors FoxO1 and FoxO3a. These effects have functional consequences *in vivo*, as neonatal SR^−/−^ mice exhibit drastic cortical overgrowth and aberrant layering. Lastly, we use unbiased proteomics to uncover putative D-cysteine-interacting proteins, an approach which identifies MARCKS as a candidate binding partner.

## RESULTS

### Distribution of D-cysteine in the mammalian brain

To search for endogenous D-cysteine in mammals, we developed assays to separate and measure L- and D-cysteine in tissue samples. High-performance liquid chromatography is the gold standard for resolution of stereoisomeric mixtures (**16**). We accordingly developed a method for labeling of cysteine enantiomers followed by chiral HPLC. Fluorescent labeling of free thiols with 4-(aminosulfonyl)-7-fluoro-2,1,3-benzoxadiazole (ABDF) (**17**) followed by enantiomeric separation using the teicoplanin-based CHIROBIOTIC T chiral column (*SI Appendix,* Fig. S1*A*) allows for robust differentiation of L- and D-cysteine in purified solution and tissue lysates (Fig. 1*A*, Fig. S1*B–C*). This novel method establishes the presence of endogenous D-cysteine in the mammalian central nervous system. D-cysteine is present at levels of 50 μM in the adult mouse brain (Fig. 1*A*), a concentration approximately one fifth that of brain D-serine (**4**). We also detect free D-cysteine in human cerebrospinal fluid at a concentration of 79 μM (Fig. S1*C*).

**Figure 1.**
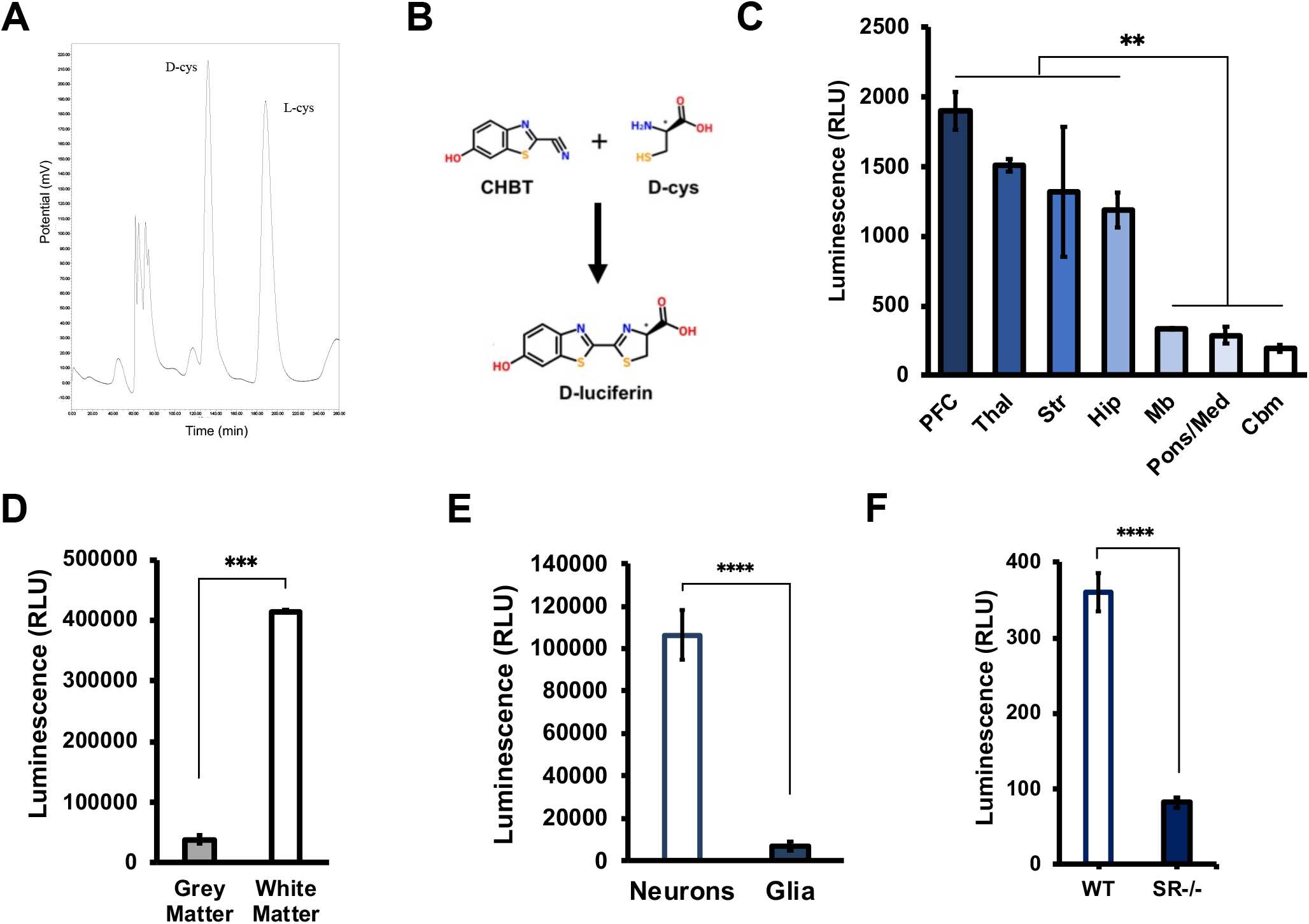
Identification of endogenous D-cysteine in the mammalian brain. (*A*) Representative chromatogram from HPLC analysis of adult mouse brain extract. (*B*) Reaction scheme for D-cysteine (D-cys) luciferase assay. Condensation of endogenous D-cys with exogenously added 2-cyano-6-hydroxybenzothiazole (CHBT) yields D-luciferin, which is oxidized by D-luciferin-specific luciferase from the firefly *P. pyralis* to produce light. (*C*) Luciferase assay measurements of relative D-cys levels in lysates of various regions of adult mouse brain (*n* = 3). *PFC*, prefrontal cortex; *Thal*, thalamus; *Str*, striatum; *Hip*, hippocampus; *Mb*, midbrain, *Pons/Med*, pons/medulla; *Cbm*, cerebellum. (*D*) Luciferase assay of D-cys levels in postmortem samples of human gray and white matter (*n* = 3). (*E*) Luciferase assay of D-cys in primary cortical neuronal and mixed glial cultures (*n* = 3). (*F*) Luciferase assay of D-cys in brain lysates from adult wild-type and serine racemase knockout (SR^−/−^) mice (*n* = 3). Data are graphed as mean ± SEM. \*\**P* < 0.01, \*\*\**P* < 0.001, \*\*\*\**P* < 0.0001, ANOVA with *post-hoc* Tukey’s test (*C*) or two-tailed unpaired student’s *t*-test (*D-F*).

While HPLC allows for sensitive determination of absolute D-cysteine concentration in tissue samples, the long elution times of this assay limit its throughput. To circumvent this issue, we took advantage of the use of D-cysteine in the industrial synthesis of D-luciferin (**18**) to develop a rapid and specific bioluminescence assay for measuring free D-cysteine. Condensation of D-cysteine with 2-cyano-6-hydroxybenzothiazole (CHBT) yields D-luciferin, which is then oxidized to oxyluciferin upon addition of ATP and luciferase from the firefly *Photinus pyralis* to generate light (Fig. 1*B*). The stereospecificity of the *P. pyralis* luciferase for D-luciferin prevents the generation of luminescence by L-cysteine. This assay readily detects low micromolar concentrations of D-cysteine (Fig. S1*D*).

We leveraged the simplicity of the luciferase assay to assess the distribution of D-cysteine in the mammalian brain. D-cysteine is enriched in regions of mouse forebrain (prefrontal cortex, thalamus, striatum, and hippocampus) compared to midbrain and hindbrain regions (pons/medulla and cerebellum), though differences between forebrain regions were not significant (Fig. 1*C*). In postmortem human brain, D-cysteine levels are tenfold higher in white matter than in grey matter (Fig. 1*D*). D-cysteine appears to be primarily localized to neurons, as its levels in cultured murine primary cortical neurons (PCNs) were 15 times greater than those found in primary mixed glial cultures from mouse cortex (Fig. 1*E*).

Cysteine and serine are structurally similar: cysteine contains a sulfhydryl group bound to its *β*-carbon, while serine contains a chemically related hydroxyl group at the same position. Accordingly, we hypothesized that SR, which generates D-serine from L-serine, might also serve as a cysteine racemase. In support of this hypothesis, we found that levels of D-cysteine are reduced by 80% in the brains of adult SR null (SR^−/−^) mice (Fig. 1*F*), suggesting that SR may serve as a biosynthetic enzyme for D-cysteine.

### Brain D-cysteine is developmentally regulated

We next interrogated potential developmental regulation of brain D-cysteine. HPLC analysis revealed that D-cysteine is dramatically enriched in the embryonic mouse brain compared to the adult (Fig. 2*A*). Brain D-cysteine concentrations are as high as 4.5 mM at embryonic day 9.5 (E9.5) and decline steadily to adult levels of 50 μM. Measurement of relative D-cysteine levels at higher temporal resolution by luciferase assay highlighted two time periods during which brain D-cysteine levels drop dramatically: during midgestation (E9.5-E13.5) and over the first two weeks of postnatal life (Fig. 2*B–C*). By contrast, D-cysteine levels are relatively stable from E13.5 to birth and during early adult life.

**Figure 2.**
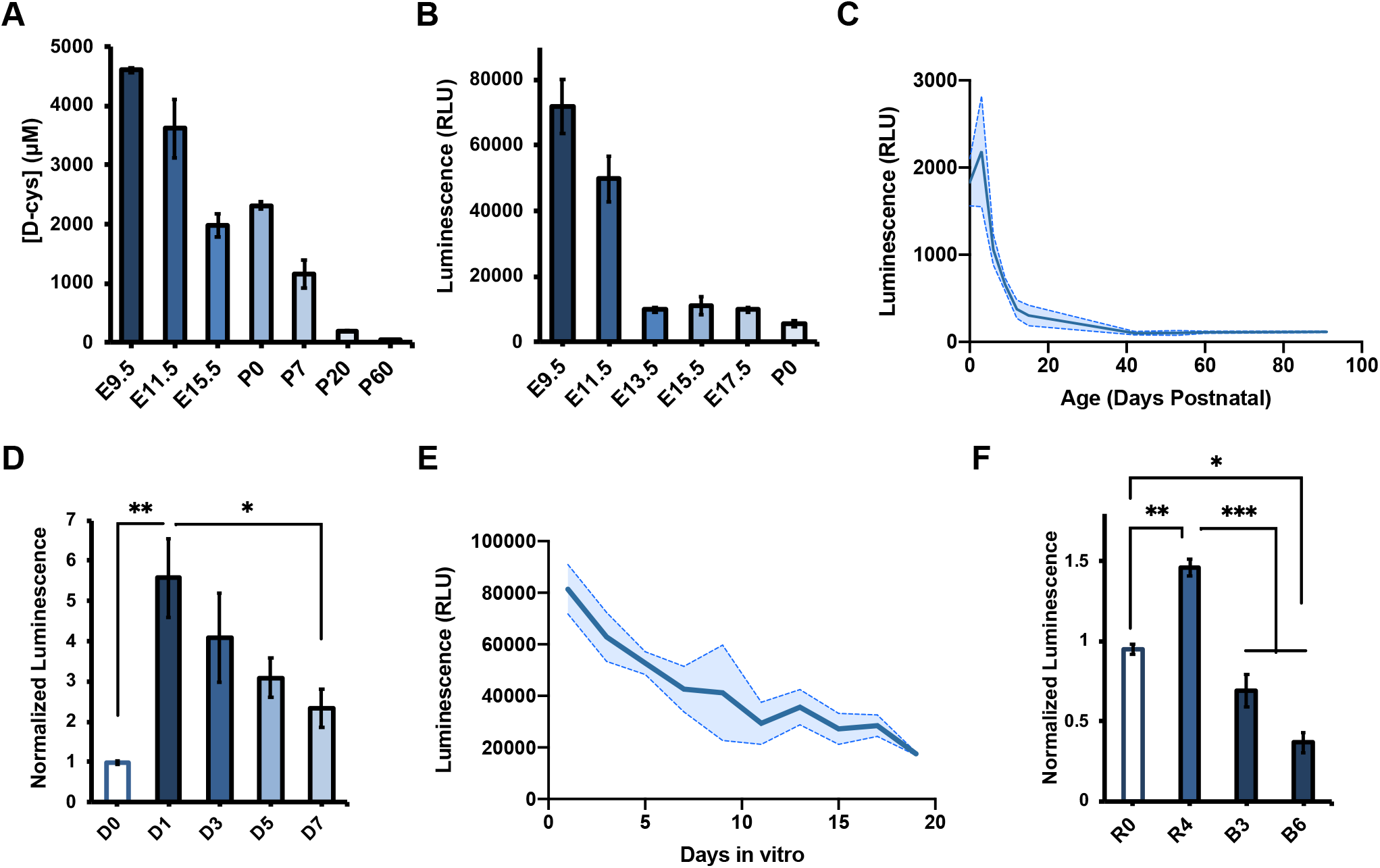
Brain D-cysteine levels are developmentally regulated. (*A-C*) D-cys levels in brain lysates from mice of indicated ages determined by (*A*) HPLC and (*B-C*) luciferase assay (*n* = 3). (*D*) Luciferase assay measurements of relative D-cys levels in primary cortical neural progenitor cells (NPCs) that were either left undifferentiated (D0) or differentiated for the indicated number of days (*n* = 3). (*E*) Luciferase assay of D-cys levels in primary cortical neurons (PCNs) cultured for the indicated number of days (*n* = 3). (*F*) Luciferase assay of D-cys levels in SK-N-SH cells left undifferentiated (R0) or differentiated with retinoic acid for 4 days (R4) followed by BDNF (B3, B6) (*n* = 3). Data are graphed as mean ± SEM. \**P* < 0.05, \*\**P* < 0.01, \*\*\**P* < 0.001, ANOVA with *post-hoc* Tukey’s test.

The two developmental periods during which D-cysteine levels decrease markedly coincide with major milestones in forebrain development. Beginning at E9, neuroepithelial stem cells, which constitute the major neural stem cell (NSC) population in the early embryonic central nervous system, begin to differentiate into radial glial cells, the primary neural progenitor cell (NPC) population of the developing brain (**19**). Similarly, radial glia and other NPCs execute their final neurogenic divisions over the course of the first week of postnatal life (**20**). These correlations suggest that D-cysteine may be involved in cell fate decisions in the developing forebrain. In order to explore this hypothesis, we isolated and cultured NPCs from E13.5-E14.5 mouse cortex and monitored cellular D-cysteine concentrations during induced differentiation into postmitotic neurons. Differentiation elicited a robust increase in D-cysteine levels 24 h after initiation of differentiation, which gradually declined over the following week (Fig. 2*D*). This decrease was mimicked in PCNs allowed to mature *in vitro* for 19 days, during which time D-cysteine levels decreased by fourfold (Fig. 2*E*).

To confirm our findings in another context and assess their potential conservation in human cells, we utilized the SK-N-SH human neuroblastoma cell line, whose subclone SH-SY5Y has been used as a model for neuronal differentiation (**21**). In this two-stage system, cells are treated with retinoic acid, which induces cell-cycle exit and initiates neurite outgrowth (i.e., differentiation phase), before treatment with brain-derived neurotrophic factor (BDNF), which further stimulates neurite outgrowth and arborization (maturation phase) (**22**). Retinoic acid-mediated differentiation increased cellular D-cysteine content by 50% after four days; induction of maturation with BDNF decreased D-cysteine levels to below untreated values after six days (Fig. 2*F*). Collectively, these data point to a role for D-cysteine in neuronal cell fate decisions.

### D-cysteine inhibits NPC proliferation through modulation of the Akt-FoxO signaling axis

Prenatal brain development reflects a balance between NPC proliferation (self-renewal) and differentiation (neurogenesis). Because NPCs are the predominant brain cell type during embryonic life, during which D-cysteine is present at high levels, we explored the influence of D-cysteine on these two fundamental properties of NPC homeostasis. Treatment with 1 mM D-cysteine, the concentration found in perinatal brain, for 48 h reduced proliferation of cultured NPCs by ~50% as determined by staining for the mitotic marker Ki-67 and the nucleotide analog 5-ethynyl-2’-deoxyuridine (EdU), which labels dividing cells (Fig. S2*A*–*B*). Importantly, these effects were not elicited by equivalent concentrations of D-serine or L-cysteine (Fig. 3*A*). Promotion of cell-cycle exit by D-cysteine was not associated with an induction of neuronal differentiation, as D-cysteine treatment did not significantly alter the numbers of newborn or mature neurons present in the cultures as monitored by doublecortin and MAP2 immunofluorescence, respectively (Fig. S2*D–E*).

**Figure 3.**
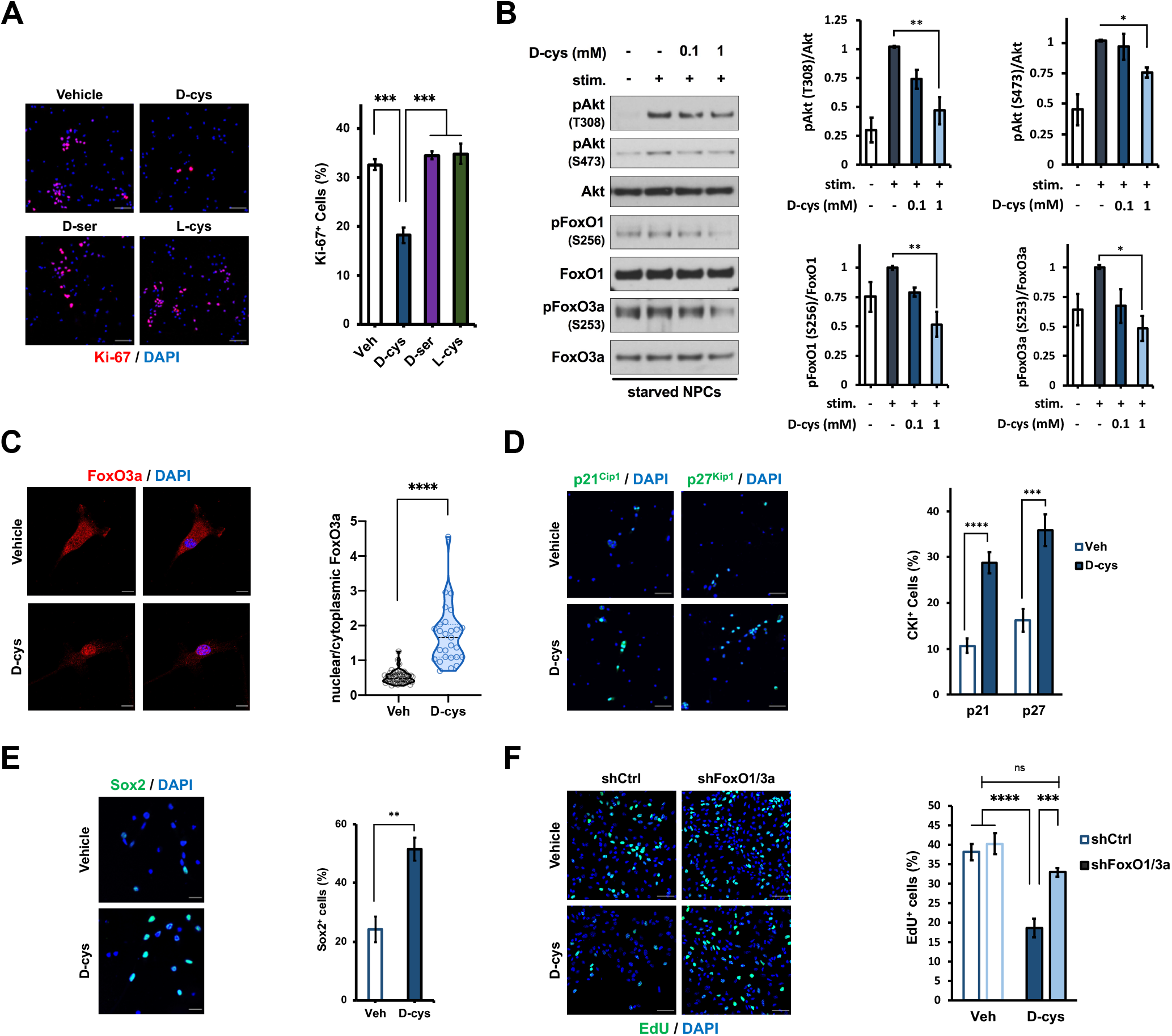
D-cysteine inhibits neural progenitor cell proliferation through activation of FoxOs. (*A*) Representative images and quantification of immunostaining for Ki-67 and DAPI in NPCs treated with saline (vehicle) or 1 mM of the indicated amino acid for 48 h (*n* = 3). Data were quantified as the number of Ki-67-positive (Ki-67^+^) cells as a percent of the total number of (DAPI^+^) cells. Scale bar 50 μm. (*B*) Representative immunoblots and quantification of phospho- and total Akt, FoxO1, and FoxO3a in NPCs. Cells were starved overnight followed by treatment with vehicle or the indicated concentration of D-cys for 1 h, followed by stimulation with complete medium for 5 minutes. Data are expressed as the normalized ratio of phosphorylated to total protein in each condition (*n* = 3). (*C*) Immunostaining for FoxO3a and DAPI in NPCs treated with vehicle or 1 mM D-cys for 48 h. Data are expressed as the ratio of the mean FoxO3a fluorescence intensity in the nucleus (DAPI-positive area) compared to the cytoplasm (*n* = 25-35 cells per group). Scale bar 10 μm. (*D-E*) Immunostaining for (*D*) p21^Cip1^, p27^Kip1^, (*E*) Sox2, and DAPI in NPCs treated with vehicle or 1 mM D-cys for 48 h. Data were quantified as in (*A*) (*n* = 3). Scale bar 50 μm (*D*) and 20 μm (*E*). (*F*) EdU and DAPI staining in NPCs. Cells were infected with lentivirus producing shRNA against a non-targeting control (shCtrl) or shRNAs targeting FoxO1 and FoxO3a for 72 h before treatment with vehicle or 1 mM D-cys for 48 h, followed by addition of 10 μM EdU for 2 h. Data were quantified as in (*A*) (*n* = 3). Scale bar 50 μm. Bar graphs depict mean ± SEM; violin plots show median (dark line) ± interquartile range (dashed lines). \**P* < 0.05, \*\**P* < 0.01, \*\*\**P* < 0.001, \*\*\*\**P* < 0.0001, ns = *P* > 0.05, ANOVA with *post-hoc* Tukey’s test (*A, B, F*) or two-tailed unpaired student’s *t*-test (*C-E*).

Kimura and associates (**23**) recently reported that exogenous D-cysteine can serve as a metabolic precursor for hydrogen sulfide (H_2_S) upon oxidation by DAAO. To determine whether D-cysteine inhibits NPC proliferation via H_2_S production, we treated NPCs with D-cysteine and the DAAO inhibitor (**24**) 6-chloro-1,2-benzisoxazol-3(2H)-one (CBIO). Consistent with DAAO functioning as a physiologic degradative enzyme for D-cysteine, CBIO treatment alone phenocopied the antiproliferative effect of D-cysteine and potentiated its influence on NPC proliferation (Fig. S2*C*). This result suggests that the antiproliferative actions of D-cysteine in NPCs are independent of H_2_S production.

Cell proliferation is regulated by myriad signaling pathways, many of which converge on protein kinase B/Akt (**25**). We therefore hypothesized that D-cysteine inhibits Akt signaling. Accordingly, one-hour D-cysteine pretreatment dose-dependently inhibited the induction of Akt phosphorylation at Thr308 and Ser473 elicited by growth factor supplement stimulation (**26**) of growth factor-starved NPCs (Fig. 3*B*). Akt can phosphorylate and thereby regulate the activity diverse proteins. We assessed the influence of D-cysteine on several of the most prominent Akt targets. The most dramatic influence of D-cysteine was on phosphorylation of the forkhead box O transcription factors FoxO1 and FoxO3a, which are negatively regulated by Akt (**27**). In accordance with the 50% reduction of NPC proliferation, 1 mM D-cysteine reduced phosphorylation of FoxO1 (at Ser256) and FoxO3a (Ser253) by approximately 50% (Fig. 3*B*). D-cysteine also reduced phosphorylation of glycogen synthase kinase 3β(GSK-3β) (**28**), albeit to a much lesser extent (Fig. S2*F*). By contrast, D-cysteine did not affect the activity of the mammalian target of rapamycin complex 1 (mTORC1) (**29, 30**) as determined by phosphorylation of its targets p70 S6 kinase (S6K) and eukaryotic translation initiation factor 4E-binding protein 1 (4E-BP1) (Fig. S2*F*). 48 h treatment with 1 mM D-cysteine was sufficient to reduce Akt, FoxO1, and FoxO3a phosphorylation in the absence of starvation or growth factor manipulations (Fig. S2*H*).

We next focused on the actions of D-cysteine upon FoxO-mediated signaling in NPCs. FoxOs are also inactivated by serum and glucocorticoid-regulated kinase 1 (SGK1) (**31**), another target of which is N-myc downstream regulated gene 1 (NDRG1) (**32**). D-cysteine reduced NDRG1 phosphorylation at Thr346 in NPCs (Fig. S2*G*), suggesting that SGK1 activity is inhibited and may thus contribute to the activation of FoxOs by D-cysteine. SGK1 and Akt are both downstream of phosphoinositide 3-kinase (PI3K); thus, D-cysteine may act upstream of Akt and influence PI3K activity to modulate NPC proliferation (**33**).

Akt-mediated phosphorylation of FoxOs promotes their binding to 14-3-3 chaperones, which sequester FoxOs in the cytoplasm and thereby inhibit their transcriptional activity (**27**). Accordingly, D-cysteine, which inhibits Akt-mediated FoxO phosphorylation, stimulates the nuclear translocation of FoxO3a (Fig. 3*C*) and induces expression of FoxO target genes including the cyclin-dependent kinase inhibitors p21^Cip1^ and p27^Kip1^, which mediate cell cycle exit (**34**), and the pluripotency factor Sox2 (**35**), which is essential for the maintenance of NPC identity (**36**) (Fig. 3*D-E*). FoxOs are also central regulators of programmed cell death (**27, 31**), a process essential for proper brain development (**37–39**). Thus, D-cysteine stimulated caspase 3 cleavage, an essential step in apoptosis (**38**), after 48 h (Fig. S2*I*). Crucially, lentiviral shRNA-mediated double knockdown of FoxO1 and FoxO3a to 28% and 40% of control levels, respectively (Fig. S2*J*), prevented the inhibition of NPC proliferation by D-cysteine (Fig. 3*F*), demonstrating that FoxO-dependent transcription is the crucial effector of D-cysteine signaling in NPCs.

### Expansion and aberrant lamination of the cerebral cortex in perinatal SR^−/−^ mice

In light of the finding that D-cysteine, but not D-serine, regulates cortical NPC proliferation, we made use of SR^−/−^ mice as a model to explore the influence of D-cysteine upon cortical development *in vivo*. We first confirmed loss of D-cysteine in the SR^−/−^ brain at developmentally relevant time points. In contrast to adult SR^−/−^ mice, whose brains contain nearly 80% less D-cysteine than wild-type controls, E16.5 and P0 SR^−/−^ brains had only 40-50% less D-cysteine than wild-type brains (Fig. 4*A*), suggesting that SR regulates brain D-cysteine levels in an age-dependent manner. However, we suspected that reducing D-cysteine levels by half would still be sufficient to induce neurodevelopmental alterations. Indeed, cortices of E16.5 SR^−/−^ mice were 32% thicker than wild-type counterparts, with expansion being particularly pronounced in neurogenic zones (cortical plate, subplate, and intermediate zone) but also evident in the subventricular zone (SVZ) (Fig. 4*B*). Proliferative NPCs reside in the subventricular and ventricular zones (VZ) of the developing cortex (**19**). Loss of D-cysteine would be expected to disinhibit NPC proliferation resulting in an expansion of the NPC population. Thus, a greater fraction of VZ/SVZ cells stained positive for the NPC marker Sox2 in E16.5 SR^−/−^ cortices (Fig. 4*C*).

**Figure 4.**
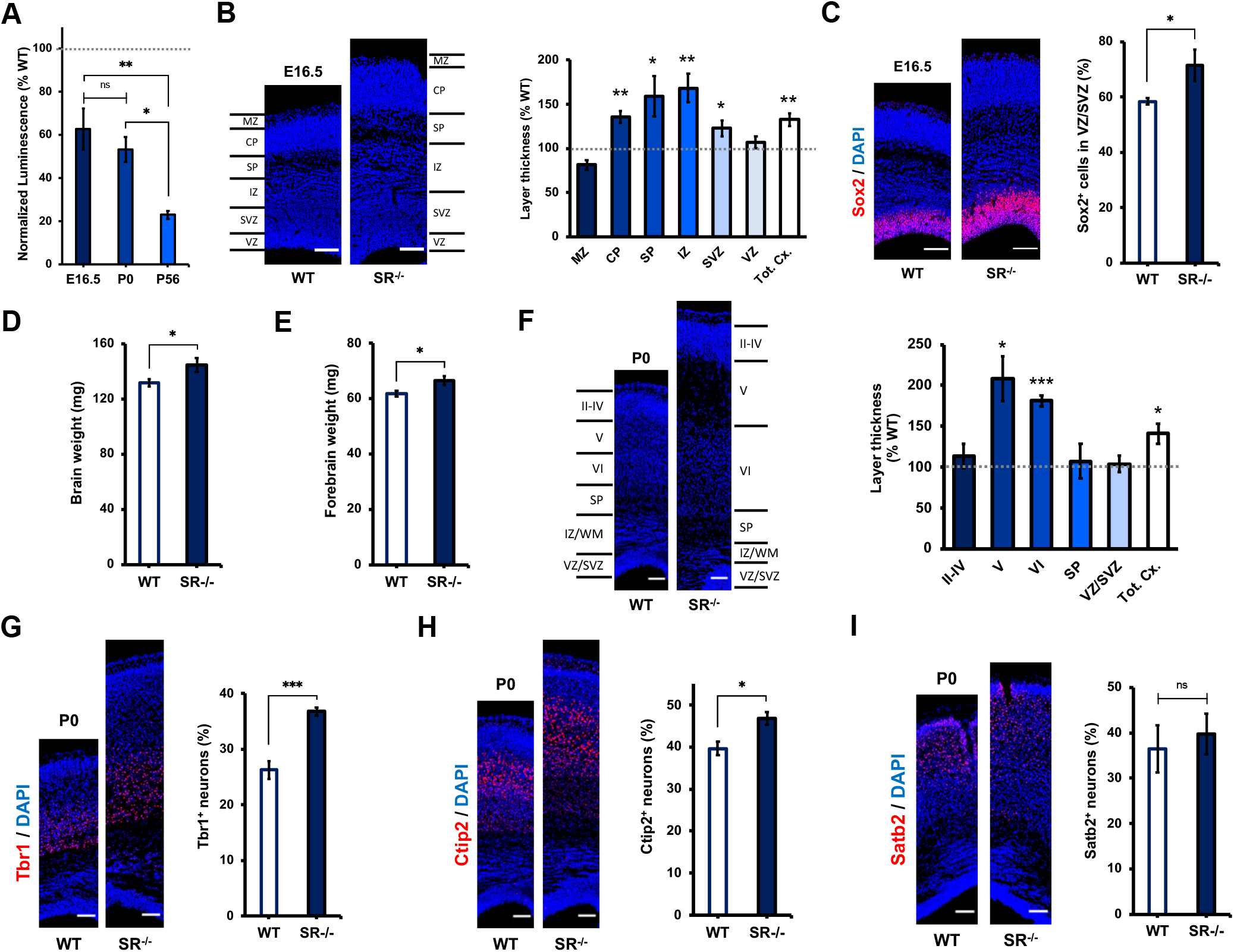
Aberrant cortical architecture in embryonic and neonatal SR^−/−^ mice. (*A*) Luciferase measurements of D-cys levels in SR^−/−^ mouse brain lysates at the indicated ages. Data are normalized to luciferase measurements in age-matched wild-type controls (*n* = 4-5) (*B*) Representative DAPI staining and quantification of cortical zone thickness in coronal cortical sections from E16.5 WT and SR^−/−^ mice. Layer thicknesses in SR^−/−^ mice are normalized to the mean thickness of the corresponding layer in WT mice (*n* = 4-5). *MZ*, marginal zone; *CP*, cortical plate; *SP*, subplate; *IZ*, intermediate zone, *SVZ*, subventricular zone; *VZ*, ventricular zone; *Tot. Cx.*, total cortical thickness. (*C*) Immunostaining for Sox2 and DAPI in E16.5 WT and SR^−/−^ cortical sections. Data are expressed as the ratio of Sox2^+^ cells to total (DAPI^+^) cells in the VZ/SVZ (*n* = 4-5) (*D-E*) Weight of (*D*) whole brains and (*E*) forebrains of P0 WT and SR^−/−^ mice (*n* = 8-11). (*F*) Representative DAPI staining and quantification of cortical layer thickness in P0 WT and SR^−/−^ mice, quantified as in (*B*) (*n* = 4-5). (*G-I*) Immunostaining for (*G*) Tbr1, (*H*) Ctip2, (*I*) Satb2, and DAPI in P0 WT and SR^−/−^ cortical sections. Data are quantified as the ratio of marker-positive cells to the total number (DAPI^+^) of cells (*n* = 4-5). Scale bars 100 μm. Data are graphed as mean ± SEM. \**P* < 0.05, \*\**P* < 0.01, \*\*\**P* < 0.001, ns = *P* > 0.05, ANOVA with *post-hoc* Tukey’s test (*A*) or two-tailed unpaired student’s *t*-test (*B-I*).

We continued our examination of developmental aberrations in neonatal (postnatal day 0, P0) SR^−/−^ mice, which exhibited increased weight of both whole brain and forebrain (Fig. 4*D–E*). Cortical thickening was even more pronounced in P0 brain, with a 41% increase in overall cortical thickness in SR^−/−^ mice. This expansion was accounted for by a near doubling of the thickness of deep cortical layers (layers V-VI), while layer II-IV thickness was unaffected (Fig. 4*F*). This expansion would most likely be due to an overproduction of early-born deep-layer neurons, which could result from an increase in the size of the progenitor pool as observed above. Accordingly, a greater percentage of SR^−/−^ cortical cells stained positive for Tbr1 and Ctip2, markers of subplate/layer VI and layer V neurons (**40, 41**), respectively (Fig. 4*G–H*). By contrast, numbers of Satb2^+^ later-born upper-layer neurons (**42**) were unchanged (Fig. 4*I*). Together, these data support the hypothesis that loss of D-cysteine promotes expansion of the cortical NPC population and thereby increases cortical size through overproduction of deep-layer neurons.

### Characterization of MARCKS as a putative D-cysteine-binding protein

In order to further clarify the molecular functions of D-cysteine, we employed an unbiased proteomic approach to screen for potential D-cysteine binding partners in NPCs. We immunoprecipitated proteins from D-cysteine-treated NPCs with either an anti-D-cysteine antibody or a control preimmune immunoglobulin, after which proteins eluted by D-cysteine were analyzed by liquid chromatography-tandem mass spectrometry (LC-MS/MS). We filtered identified proteins by their frequency of occurrence in the contaminant repository for affinity purification (CRAPome) (**43**). We noted an increase in immunoprecipitated proteins with CRAPome frequency ≥0.4; accordingly, we prioritized proteins below this threshold as putative true-positive hits (Fig. S3*A*). 44 proteins met this criterion, among which were the myristoylated alanine-rich C-kinase substrate (MARCKS) and its homolog MARCKSL1 (Fig. 5*A*, Fig. S3*B–D*). We were particularly intrigued by the presence of MARCKS, as it is known to interface with PI3K signaling (**44**).

**Figure 5.**
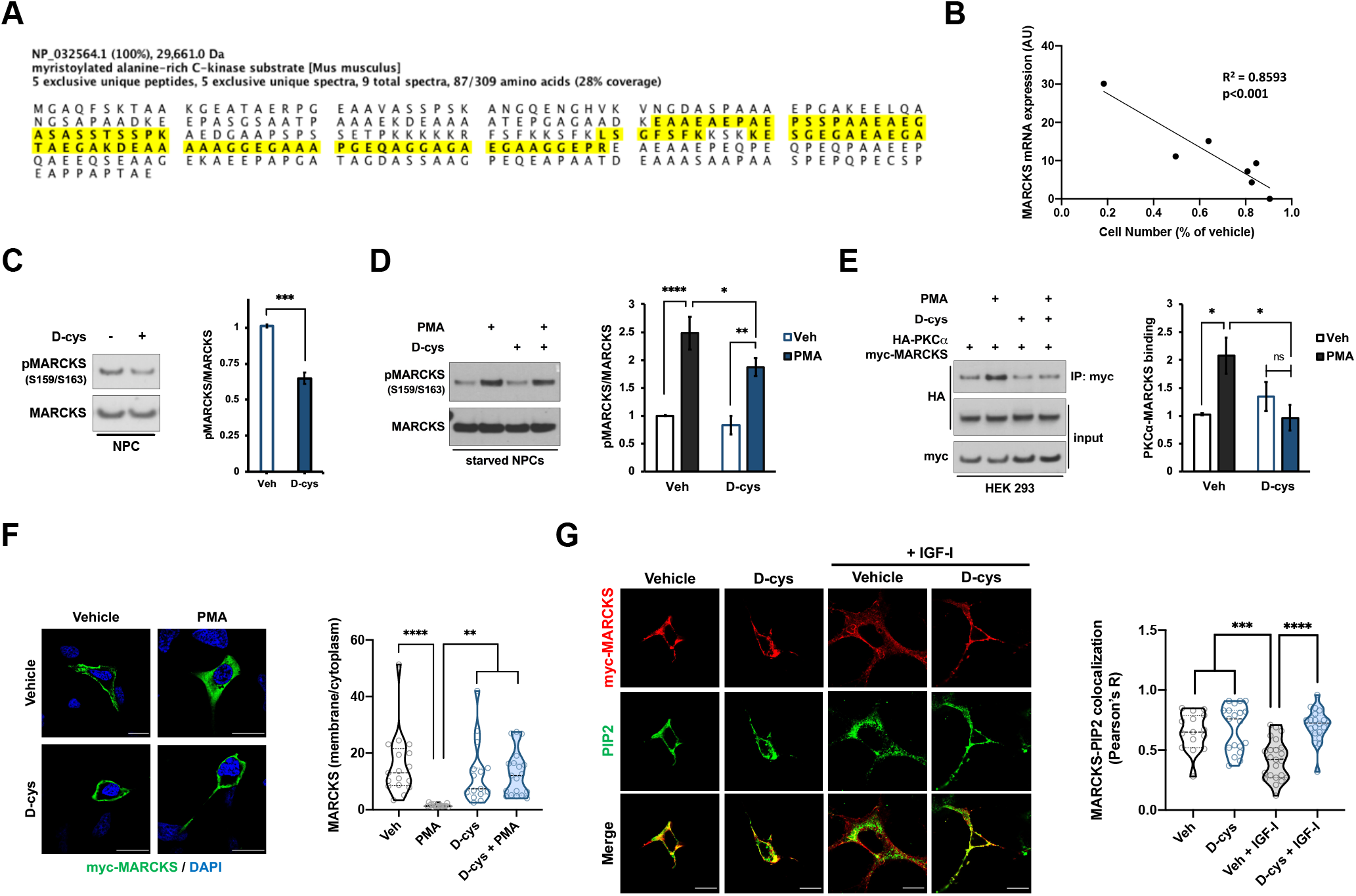
MARCKS is a putative D-cysteine-interacting protein. (*A*) Amino acid sequence of mouse MARCKS protein. Peptides identified by mass spectrometric analysis of anti-D-cys immunoprecipitates are highlighted in yellow. (*B*) Correlation between MARCKS mRNA expression (as determined by the Human Protein Atlas (**80**), available at http://www.proteinatlas.org) and antiproliferative effect of D-cys, as determined by the ratio of cell number after treatment with 1 mM D-cys for 48 h compared with vehicle-treated cells, in a panel of human cancer cell lines (*n* = 3). (*C-D*) Immunoblots and quantification of MARCKS phosphorylation in NPCs that were either (*C*) treated with vehicle or 1 mM D-cys for 48 h or (*D*) starved overnight followed by 20 min treatment with vehicle, 1 mM D-cys, and/or 250 nM phorbol 12-myristate 13-acetate (PMA) as indicated (*n* = 4). (*E*) Immunoblots and quantification of lysates (input) and anti-myc immunoprecipitates (IP) from HEK 293 cells overexpressing HA-PKC*α* and myc-MARCKS constructs. Cells were starved overnight followed by 20 min treatment with vehicle, 1 mM D-cys, and/or 500 nM PMA as indicated. Data are expressed as ratio of HA-PKC*α* in the IP fraction to HA-PKC*α* in the input fraction (*n* = 3). (*F*) Representative images and quantification of immunostaining for myc and DAPI in HEK 293 cells overexpressing myc-MARCKS and treated as in (*E*). Data are expressed as the ratio of the mean myc-MARCKS fluorescence intensity in the membrane compared to the cytoplasm (*n* = 15-20 cells per group). Scale bar 20 μm. (*G*) Immunostaining for myc and PIP2 in HEK 293 cells overexpressing myc-MARCKS. Cells were starved overnight before treatment with vehicle or 1 mM D-cys for 1 h followed by treatment with 100 ng/mL IGF-I for 10 min. Data are expressed as the Pearson correlation coefficient between myc and PIP2 stains at the plasma membrane (*n* = 3). Scale bar 15 μm. Bar graphs depict mean ± SEM; violin plots show median (dark line) ± interquartile range (dashed lines). \**P* < 0.05, \*\**P* < 0.01, \*\*\**P* < 0.001, \*\*\*\**P* < 0.0001, ns = *P* > 0.05, sum of squares *F*-test (*B*), two-tailed unpaired student’s *t*-test (*C*), or ANOVA with *post-hoc* Tukey’s test (*D-G*).

Basic bioinformatic analyses of published datasets (**45, 46**) revealed that *Marcks* mRNA is enriched in progenitor compared to neuronal cell populations in E14.5 mouse brain (Fig. S3*F*). Like D-cysteine, *MARCKS* mRNA expression decreases in the brain over the course of embryonic and postnatal development (Fig. S3*G*). By contrast, transcript expression of the NMDAR GluN1 subunit, which acts as the receptor for D-serine, is enriched in neuronal populations and increases over developmental time (Fig. S3*F–G*). *Srr* mRNA is also enriched in neuronal populations, consistent with the greater loss of D-cysteine in adult versus perinatal SR^−/−^ mice (Fig. S3*F*). Importantly, *MARCKS* mRNA expression was strongly correlated (R^2^ = 0.8593, *P* < 0.001) with the magnitude of proliferation inhibition by D-cysteine in a panel of human cancer cell lines (Fig. 5*B*, Fig. S3*E*). These data indicate that MARCKS is present in the cell types upon which D-cysteine exerts its actions and displays a developmental expression profile consistent with that of D-cysteine, supporting the notion that it may be a physiologically relevant D-cysteine-interacting protein.

We next interrogated the impact of D-cysteine on MARCKS-mediated signaling. 48 h treatment with 1 mM D-cysteine reduced MARCKS phosphorylation at Ser159/163 in NPCs (Fig. 5*C*). As its name suggests, MARCKS is the major substrate of protein kinase C (PKC) (**47**). Upon activation by calcium, diacylglycerol, or other stimuli, PKC binds and phosphorylates MARCKS, an action which translocates MARCKS from the plasma membrane, where it is basally localized, into the cytoplasm (**48**). D-cysteine reduced phosphorylation of MARCKS elicited by the PKC activator phorbol 12-myristate 13-acetate (PMA) in starved NPCs (Fig. 5*D*) and prevented the binding of overexpressed myc-tagged MARCKS and HA-tagged PKCα induced by PMA in starved human embryonic kidney (HEK 293) cells (Fig. 5*E*). D-cysteine further prevented the ability of PMA to stimulate myc-MARCKS internalization in HEK 293 cells (Fig. 5*F*).

MARCKS is known to associate with the lipid phosphatidylinositol 4,5-bisphosphate (PIP2) at the membrane during inactive (starved) cellular conditions (**47**). Upon growth factor stimulation, activation of MARCKS by PKC and other kinases dissociates it from PIP2, freeing PIP2 to be phosphorylated by PI3K to form phosphatidylinositol 3,4,5-bisphosphate (PIP3), the essential molecule for Akt activation (**49**). Stimulation of starved HEK 293 cells with insulin-like growth factor I (IGF-I) decreases the colocalization of PIP2 and myc-MARCKS at the membrane, and D-cysteine completely blocks this decrease (Fig. 5*G*). Thus, D-cysteine causes MARCKS to sequester PIP2 away from PI3K in response to growth factor signaling, providing a mechanistic basis for inhibition of Akt signaling by D-cysteine. While preliminary, these findings suggest that MARCKS may be a physiologic interactor of D-cysteine.

## DISCUSSION

D-amino acids, once thought to be “unnatural” isomers, are an important class of signaling molecules whose roles in mammalian neurobiology continue to be unearthed. Our present work describes the newest member of this family, D-cysteine. By developing sensitive and specific assays for the detection of D-cysteine, we report for the first time the identification of endogenous D-cysteine in mammals. We delineate its actions in cortical development and identify a signaling axis and putative binding partner mediating these effects. Our findings not only have implications for the understanding of the functions of D-amino acids in the nervous system, but also provide insight into the regulation of NPC homeostasis in the developing brain by small molecules.

The regional distribution of D-cysteine in the adult brain closely parallels that of SR and D-serine (**9, 50**), with enrichment in forebrain regions including the prefrontal cortex and hippocampus. Conversely, D-cysteine localizations are reciprocal to those of DAAO (**50**). These data are consistent with conceptualizations of SR and DAAO as biosynthetic and degradative enzymes for D-cysteine, respectively (**23**). It will be of interest to explore the extent to which modulation of D-cysteine levels contributes to previously observed phenotypes of SR^−/−^ and DAAO^−/−^ mice.

In contrast to their similar regional localizations, the temporal distributions of D-cysteine and D-serine are inversely related. Brain D-cysteine content drops drastically during early postnatal life, while D-serine levels increase over the same period (**51**). Interestingly, D-aspartate shows a similar developmental expression profile as D-cysteine (**2, 51, 52**). SR expression parallels that of D-serine in mouse brain (**51**), though its transcript expression, along with D-serine levels (**52**), is mostly stable during prenatal development in human brain (Fig. S3*G*). SR is further enriched in neuronal compared to progenitor populations in the embryonic mouse brain (Fig. S3*F*), and while adult SR^−/−^ mice lack 80% of brain D-cysteine, D-cysteine levels are decreased by only 40% and 50% in embryonic and neonatal SR^−/−^ brains, respectively. Collectively, these observations imply the existence of SR-independent pathways for D-cysteine production in the developing brain.

Our analyses of D-cysteine actions in NPCs suggest its major function is to inhibit their proliferation. This finding appears somewhat contradictory, as D-cysteine is present at high levels during developmental periods in which NSC/NPC proliferation is widespread. However, as proliferative growth factors are present at high levels at these time points (**19, 20, 53**), the existence of endogenous inhibitors or “brakes” on growth factor signaling serves a homeostatic function, as evidenced by macrocephaly and neurodevelopmental disorders in patients and animal models containing mutations in negative regulators of growth factor signaling cascades (**54–56**). By inactivating Akt, D-cysteine may serve as such a brake, in a manner similar to the phosphatase and tensin homolog (PTEN) and other inhibitors of growth factor/Akt signaling (**57, 58**). Our findings support a model wherein, as the fraction of proliferative cells decreases over development due to an increase in terminally differentiated cells (neurons), the proliferation-limiting effects of D-cysteine become progressively less necessary and its production is decreased.

We identified the transcription factors FoxO1 and FoxO3a as the crucial effectors of D-cysteine in NPCs. FoxOs are established negative regulators of NPC proliferation and neurogenesis (**59, 60**). While we did observe increased numbers of deep-layer cortical neurons in SR^−/−^ mice, apparently due to an expanded NPC pool, D-cysteine did not have a direct effect on neurogenic differentiation of NPCs *in vitro*. However, by activating FoxOs, D-cysteine may inhibit differentiation in response to extrinsic signals, particularly by inducing expression of Sox2, which opposes neurogenesis (**36**). FoxOs and Sox2 specify the stem cell identity of populations including NPCs (**35, 36, 59, 60**). Accordingly, D-cysteine may itself function to maintain NSC/NPC identity. This hypothesis aligns well with our finding that D-cysteine levels decrease rapidly from E9.5-E13.5 and P0-P7, two periods of widespread NSC/NPC differentiation (**19, 20**). Importantly, further work will be required to determine whether D-cysteine acts on NPCs in a paracrine or autocrine/cell-autonomous manner *in vivo*.

We found that D-cysteine, but not D-serine, reduces proliferation of NPCs. Thus, the neurodevelopmental aberrations observed in SR^−/−^ mice, which stem from expansion of the NPC population, appear to be attributable to D-cysteine. However, our data do not exclude a role for D-serine in related cellular events during corticogenesis.

Our proteomic analysis of anti-D-cysteine immunocomplexes identified MARCKS as a putative D-cysteine interactor. MARCKS has established roles in forebrain development, and its germline knockout is embryonic lethal in mice (**61**). MARCKS^−/−^ mice display aberrant cortical lamination and incomplete fusion of the cerebral hemispheres as well as area-specific defects in NPC proliferation and migration (**62**). Sequestration of PIP2 by MARCKS regulates PI3K-dependent signaling (**49**) and thus provides a mechanistic link between D-cysteine-MARCKS interactions and D-cysteine-mediated Akt inhibition. We previously demonstrated that SR is physiologically inhibited by PIP2 (**63**), suggesting the potential existence of an autoregulatory feedback loop for D-cysteine production and signaling through MARCKS. Future studies will be required to delineate the consequences of D-cysteine-MARCKS binding on cortical development *in vivo*.

We considered the possibility that the NMDAR might serve as a D-cysteine receptor given the structural similarity between D-cysteine and the NMDAR co-agonist D-serine (**5**). However, we did not identify any NMDAR subunits in our anti-D-cysteine immunoprecipitates, and the developmental expression profile of the D-serine-binding GluN1 subunit (**64**) is reciprocal to that of D-cysteine (Fig. S3*F-G*). Our data do not exclude the possibility of NMDAR-mediated functions of D-cysteine in the adult brain. However, in addition to being present at much lower levels than D-serine (50 μM compared to 250 μM), D-cysteine exhibits twentyfold lower affinity for forebrain synaptic NMDAR sites (**65**). H_2_S is known to facilitate NMDAR neurotransmission (**66**). Accordingly, D-cysteine may also influence NMDAR function indirectly through DAAO-mediated production of H_2_S (**23**), as D-cysteine has been shown to exert H_2_S-dependent dendritogenic effects in cultured cerebellar neurons (**67**).

The strongest link between D-amino acids and human neuropsychiatric phenotypes comes from schizophrenia. Polymorphisms in the *SRR* gene, as well as decreased D-serine levels, have been identified in multiple cohorts of schizophrenic patients (**68–71**), and SR^−/−^ mice have been found to recapitulate many of the molecular and behavioral correlates of schizophrenia (**72**). Though these phenotypes have primarily been attributed to impaired D-serine-mediated NMDAR neurotransmission (**73**), schizophrenia is increasingly appreciated as a neurodevelopmental disorder (**74–77**). Alterations in MARCKS expression in schizophrenia have also been described (**78, 79**). Accordingly, it will be of great interest to determine D-cysteine levels in schizophrenic patients and to explore the extent to which D-cysteine-dependent neurodevelopmental aberrations underlie schizophrenia-related phenotypes of SR^−/−^ mice.

## MATERIALS AND METHODS

### Reagents

D-cysteine was purchased from Santa Cruz Biotechnology. D-serine, L-cysteine, all-*trans* retinoic acid, and 6-chloro-1,2-benzisoxazol-3(2H)-one (CBIO) were purchased from Sigma Aldrich. Phorbol 12-myristate 13-acetate was purchased from Cayman Chemical. Recombinant human BDNF and IGF-I were purchased from PeproTech.

### Animals

All experiments involving animals were performed in accordance with the Johns Hopkins Medical Institutions Animal Care and Use Committee policies, as well as with NIH guidelines for use of experimental animals. C57Bl6/J mice were purchased from the Jackson Laboratory. CD-1 mice were purchased from Charles River Laboratories. Serine racemase knockout mice were generated as previously described (**81**). C57Bl6/J mice were used as wild-type controls for SR^−/−^ mice; CD-1 mice were used in all other experiments.

### Human tissue and cerebrospinal fluid samples

All work involving patients was performed in accordance with protocols approved by the Institutional Review Board at the Johns Hopkins University School of Medicine. Human brain samples from normal (non-diseased) subjects were obtained from the Rocky Mountain Multiple Sclerosis Center tissue bank. Punch samples were taken from frozen post-mortem brain slices and kept at −80 °C. Cerebrospinal fluid (CSF) was collected from patients undergoing posterior fossa craniectomies, burr holes, or lumbar punctures. CSF was centrifuged for 5 min at 2000 *g* to clear cellular material. The supernatant was then stored at −80 °C until use.

### Cell culture

#### Mouse primary cell culture

Primary cortical neurons were isolated from E15.5-E18.5 pregnant CD-1 mice as previously described (**82**) and were cultured on poly-D-lysine and poly-L-ornithine (100 μg/mL)-coated plates in Neurobasal-A medium containing antioxidant-free B27 supplement (Gibco) and 2 mM L-glutamine. Primary neural progenitor cells (NPCs) were isolated from E13.5 pregnant CD-1 mice and cultured on poly-D-lysine (20 μg/mL) and laminin (10 μg/mL)-coated plates in NeuroCult Basal Medium containing NeuroCult Proliferation Supplement (StemCell Technologies) and 20 ng/mL EGF. NPCs were differentiated by culturing in NeuroCult Basal Medium with NeuroCult Differentiation Supplement (StemCell). Primary mixed glial cultures were isolated from P3-P6 CD-1 mouse pups as previously described (**83**).

#### Immortalized cell lines

Cell lines were obtained from ATCC and cultured in the indicated base media containing 10% fetal bovine serum, 2 mM L-glutamine, and 1% penicillin/streptomycin. HEK 293, HEK 293T, and MCF-7 cells were cultured in DMEM (Corning Cellgro). SK-N-SH, U-87 MG, SK-MEL-2, Caco-2, HeLa, and HepG2 cells were cultured in EMEM (Sigma).

#### SK-N-SH differentiation

SK-N-SH cells were differentiated as previously described (**22**) with modifications. Briefly, cells were incubated in Neurobasal-A medium containing B27 supplement, 2 mM L-glutamine, and 10 μM retinoic acid for four days, at which point medium was removed and replaced with Neurobasal-A medium containing B27, 2 mM L-glutamine, and 50 ng/mL recombinant human BDNF for up to six days.

#### Starvation

HEK 293 cells were washed 4x in warm phosphate-buffered saline (PBS) and incubated overnight in serum-free DMEM (containing 2 mM L-glutamine and 1% pen/strep). Primary NPCs were starved by removing 80% of the medium and replacing with an equal volume of NeuroCult Basal Medium without any supplements.

### Plasmids

Plasmid encoding myc-tagged mouse MARCKS was purchased from Origene. Plasmid encoding HA-PKCα was a gift from Bernard Weinstein (Addgene plasmid #21232). Lentiviral plasmids encoding psPAX2 (Addgene #12260) and pMD2.G (Addgene #12259) were gifts from Didier Trono. shRNA constructs targeting FoxO1 (5’-CCGGTGGAAACCAGCCAGCTATAAACTCGAGTTTATAGCTGGCTGGTTTCCATTTTTG-3’), FoxO3a (5’-CCGGCAGCCGTGCCTTGTCAAATTCCTCGAGGAATTTGACAAGGCACGGCTGTTTTTG-3’), or non-targeting control in the pLKO.1 vector were obtained from the Mission shRNA Library (Sigma).

### Transient transfection

HEK 293 cells were transfected using PolyFect Transfection Reagent (Qiagen) according to manufacturer’s instructions. Media was changed 4 hours after transfection and cells were used for experiments 48-72 hours later.

### Lentivirus Production

HEK 293T cells were transfected with lentiviral constructs by calcium phosphate precipitation using a second-generation packaging system. Cells were transfected at a 2:1:1 ratio with plasmids encoding lentiviral shRNAs, the packaging plasmid psPAX2, and the envelope plasmid pMD2.G. The morning after transfection, the culture media was replaced, and after another 24 hours the culture media was collected and centrifuged at 2,000 *g* for 30 min at 4 °C. The supernatant was passed through low protein binding 0.45 μm filter and lentiviral particles were collected by centrifugation at 100,000 *g* for 2 hours at 4 °C. The supernatant was aspirated and the viral pellet was resuspended in DMEM with 10% FBS before addition to NPC culture media.

### Separation of L- and D-cysteine by high-performance liquid chromatography

L- and D-cysteine from tissue lysates were separated using the CHIROBIOTIC T chiral HPLC column (5 μm particle size, L × 1.D. 25 cm × 4.6 mm) (Supelco) attached to a Waters 2690 alliance separations module connected to a fluorescence detector. Solvent was 20 mM ammonium acetate (80 % v/v)-methanol (20 % v/v) followed by isocratic elution. The flow rate was 0.05 ml/min. The stereoisomers of cysteine were detected using fluorescence at excitation λ 380 and emission λ 510 nm. The column was maintained at room temperature. D-cysteine from brain lysates was extracted by homogenization in 100 mM HEPES buffer pH 7.5 using a handheld homogenizer on ice with 10-15 strokes. The homogenates were then lysed using a Branson sonicator on ice with 6 pulses of 10-15 sec each. The samples were then centrifuged at 16,000 g for 30 min at 4 °C. After centrifugation the supernatant was harvested in a pre-chilled tube. Proteins in the supernatant were precipitated by adding 12 N HCl to the lysate to a final concentration of 2 N. The sample was cooled on ice for 30 min and centrifuged at 16,000 g for 30 min at 4°C. The supernatant containing cysteine was harvested. The supernatant was neutralized by adding one volume 10 N NaOH (v/v) followed by addition of 1 volume of 1 mM 4-(aminosulfonyl)-7-fluoro-2,1,3-benzoxadiazole (ABDF) (v/v) in 200 mM sodium borate buffer pH 8.0 containing 1 mM EDTA and 0.1 volume (v/v) of 10 % tri-*n*-butylphosphine in acetonitrile to label the free thiol of cysteine. The mixture was incubated at 50 °C for 5 minutes and vortexed thoroughly after incubation. The mixture was placed on ice and 0.01 volume (v/v) of 2.5 N HCl added to neutralize the sample. The sample was then diluted 1:1 with HPLC-grade water and filtered using a Millex LG 0.2 μm syringe filter (Millipore) into an amber HPLC vial. The fluorescently labeled sample was used for injection (10 μL volume) on the pre-equilibrated chiral HPLC column and the labeled thiols detected by fluorescence. The areas under the respective peaks were quantified based on a standard curve of purified L- and D-cysteine standards.

### D-cysteine luciferase assay

Tissue samples were homogenized on ice in five volumes of lysis buffer (500 mM tricine, 100 mM MgSO_4_, 2 mM EDTA, 1% Triton X-100, pH 7.8) containing 10 mM TCEP, 200 mM K_2_CO_3_, and 0.1 mg/mL 2-cyano-6-hydroxybenzothiazole (Santa Cruz). In the case of cultured cells, culture media was aspirated, cells were washed twice with ice-cold PBS, and cells were harvested on ice in the above solution at a volume of 250 μL per 35 mm dish. Following sonication, samples were incubated at 30 °C for 10 min with shaking to allow for D-luciferin formation. Samples were then cooled on ice and cleared by centrifugation at 16,000 *g* at 4 °C. Supernatants were transferred to fresh tubes and neutralized with concentrated HCl to a pH of 7.8. ATP was added to a final concentration of 5 mM, after which D-luciferin-specific Quantilum Recombinant Luciferase (Promega) was added to a final concentration of 25 μg/mL. Samples were transferred to opaque 96-well plates and incubated at room temperature in the dark for 5 min, at which point luminescence was measured on a SpectraMax M3 microplate reader (Molecular Devices) with a 500 ms integration time. Luminescence values were normalized to total protein concentration, obtained from supernatants using colorimetric protein assay (Bio-Rad).

### Immunocytochemistry

Cells cultured on 35 mm glass-bottom microscopy dishes (MatTek) were fixed in 4% paraformaldehyde (PFA) in PBS containing 5 mM MgCl_2_, 10 mM EGTA, and 4% sucrose, then permeabilized in 0.2% Triton X-100 in TBS. Cells were blocked in TBS containing 5% donkey serum and 5% goat serum for one hour, followed by overnight incubation at 4 °C in primary antibody (listed in Table S1), diluted according to manufacturer instructions in wash buffer (20% blocking buffer in TBS). The next day, cells were washed four times in wash buffer, incubated with fluorescent secondary antibodies at for 1-2 hours at room temperature at a dilution of 1:500 in wash buffer, washed four more times, and counterstained with 10 μg/mL DAPI (Millipore). Cells were covered with ProLong Diamond Antifade Mountant (ThermoFisher Scientific) and a glass coverslip. Images were acquired on Zeiss LSM 700 and LSM 800 confocal microscopes with ZEN Blue 2.3 software (Zeiss).

### EdU Incorporation Assay

Cells were treated with 10 μM 5-ethynyl-2’-deoxyuridine [EdU] (Cayman) in culture media for 3 hours at 37 °C, chased with 3 washes of warm PBS, then fixed and permeabilized as described above. Cells were then incubated in staining solution consisting of 100 mM Tris pH 8.5, 2 mM CuSO_4_, 10 μM AFDye 488 Azide (Click Chemistry Tools), and 100 mM ascorbic acid in the dark at room temperature for 30 minutes. Cells were then counterstained with DAPI and imaged as above.

### Immunohistochemistry

Brains from embryonic or neonatal C57Bl6/J and SR^−/−^ mice were dissected out and incubated in 4% PFA in PBS for 24-48 hours at 4 °C followed by incubation for 24-48 hours at 4 °C in 30% sucrose in PBS. Brains were then embedded in OCT Compound (Fisher) and sectioned at 30 μm thickness on a Microm Sliding Microtome. Sections were mounted to slides and permeabilized in 0.5% Triton X-100 (TX-100) in Tris-buffered saline (TBS) for 15 minutes followed by blocking in TBS+ (TBS containing 0.05% TX-100) with 5% donkey serum and 5% goat serum for one hour at room temperature. Sections were incubated overnight at 4 °C in primary antibody, diluted according to manufacturer instructions in wash buffer (20% blocking buffer in TBS+). The following day, slides were washed four times in TBS+, incubated with fluorescent secondary antibodies (1:500 dilution in wash buffer) for 2 hours at room temperature, counterstained with 10 μg/mL DAPI, and washed four times in TBS+, and covered with Vectashield Plus Antifade mounting medium (Vector Labs) and a glass coverslip. Cortical sections were imaged at equivalent points along the anterior-posterior axis in all mice.

### Western blot

Cultured cells were washed with ice-cold PBS and harvested on ice in RIPA buffer (Cell Signaling Technology) containing 50 mM sodium fluoride, 400 μM sodium orthovanadate, and protease inhibitor cocktail (Sigma). Lysates were sonicated on ice and centrifuged at 1,000 *g* for 10 min at 4 °C. β-mercaptoethanol was added to supernatants to a final concentration of 10%, and samples were mixed with LDS Sample Buffer (ThermoFisher) and boiled for 5 min. 20 μg of protein was resolved by SDS-PAGE. Protein was transferred onto PVDF membranes, which were blocked in 5% bovine serum albumin in TBS with 0.1% Tween-20 (TBST) followed by overnight incubation in primary antibodies (diluted 1:1000 in blocking buffer) at 4 °C. Membranes were washed in TBST and incubated in HRP-tagged secondary antibodies (diluted 1:5000 in blocking buffer) for one hour at room temperature. Proteins were visualized using SuperSignal West Pico Plus Chemiluminescent Substrate (ThermoFisher) and analyzed using ImageJ software.

### Immunoprecipitation

Treated cells were harvested on ice in lysis buffer (50 mM Tris HCl, 150 mM NaCl, 1% Triton X-100, and 5% glycerol (v/v)) supplemented with 50 mM sodium fluoride, 400 μM sodium orthovanadate, and protease inhibitor cocktail (Sigma). Lysates were passed 10 times through a 27G syringe and centrifuged at 10,000 *g* for 10 min at 4 °C. Lysates were then incubated with either anti-c-myc beads (Pierce) or anti-D-cys antibody overnight at 4 °C. The following day, Protein G affinity beads (Sigma) were added to the lysate (for anti-D-cys immunoprecipitation only) and incubated at 4 °C for 2 hours. Beads were then pelleted by centrifugation at 1,000 *g* for 1 min at 4 °C. The supernatant was aspirated and beads were washed 5 times in lysis buffer (or TBS for anti-D-cys IP) by centrifugation, aspiration, and resuspension. Epitope-tagged proteins were eluted from beads by boiling for 5 min in LDS sample buffer (ThermoFisher). Anti-D-cysteine immunoprecipitates were eluted by incubating beads in 1 mM D-cys in TBS for 30 min at room temperature. Eluates were then sent for mass spectrometric analysis or analyzed by Western blot as described above.

### LC-MS/MS analysis

Liquid chromatography-tandem mass spectrometry (LC-MS/MS) analysis was performed as described in (**84**) with modifications. Proteins eluted after immunoprecipitation were adjusted to pH 8.0, reduced with 7.5 mg/mL dithiothreitol at 60 ° C for 1 h and alkylated with 18.5 mg/mL iodoacetamide at RT for 15 min, and proteolyzed with 20 ng/μL trypsin in Tris/triethylammonium bicarbonate buffer containing 5% acetonitrile (ACN) overnight at 37 °C. The resultant peptides were desalted on Oasis u-HLB plates (Waters), eluted with 60% ACN/0.1% trifluoroacetic acid, and dried. Peptides were then analyzed by nano-LC-MS/MS on nano-LC-QExative Plus (ThermoFisher) in FTFT interfaced with the nano-Acquity LC system (Waters) by reverse-phase chromatography with a 2%-90% acetonitrile/0.1% formic acid gradient. Precursor ions were fragmented by higher-energy collisional dissociation with an activation energy of 27-28. Precursor and fragment ions were analyzed with the following parameters, respectively: resolution: 140000 and 53000, isolation at 1.2 and 0.8 Da, max IT of 250 and 200 ms, dynamic exclusion time 10 sec and 15 sec. MS/MS spectra were processed with Proteome Discoverer (v2.3, ThermoFisher) and queried with Mascot (v2.6.2, Matrix Science) using the following specifications: enzyme, trypsin; missed cleavage 2; precursor mass tolerance 5 ppm; fragment mass tolerance 0.011 Da; variable modifications, carbamidomethyl (C), oxidation (M), deamidation (NQ). Identified proteins/peptides were analyzed/visualized using Scaffold (Proteome Software).

### Data analysis

Analysis of fluorescence microscopy and Western blot images was performed using ImageJ software (NIH). Analysis of membrane localization in immunofluorescence experiments was performed as described (**85**), and colocalization analysis was performed using the ImageJ Coloc 2 plugin. Statistical analysis was performed in Prism (Graphpad), and graphs were generated in Prism and Excel (Microsoft). For *in vivo* experiments, *n* represents the number of mice per group/genotype; for *in vitro* experiments, *n* represents the number of independent biological replicates, unless otherwise indicated.

### Data availability

All study data are included in the article and/or supporting information.

## ACKNOWLEDGMENTS

We are grateful to Roxanne Barrow, Lauren Albacarys, Adele Snowman, Lynda Hester, and Susan McTeer (S.H.S laboratory) for their assistance, Tatiana Boronina and Bob Cole (Johns Hopkins School of Medicine Mass Spectrometry and Proteomics Core) for assistance with mass spectrometry experiments, and members of the S.H.S. laboratory for helpful discussion. We thank the Rocky Mountain Multiple Sclerosis Center for supplying post-mortem human brain tissue samples. This work was supported by US Public Health Service grant P50DA044123.

## AUTHOR CONTRIBUTIONS

E.R.S., M.M.H, S.H.S, and R.R. designed research, E.R.S, M.M.H, A.P.M, E.A., and R.R. performed research, C.V, M.M.G, and M.D.K. contributed new reagents/analytic tools, and E.R.S, M.M.H, S.H.S, and R.R. analyzed data and wrote the paper.

---

**Figure S1.**
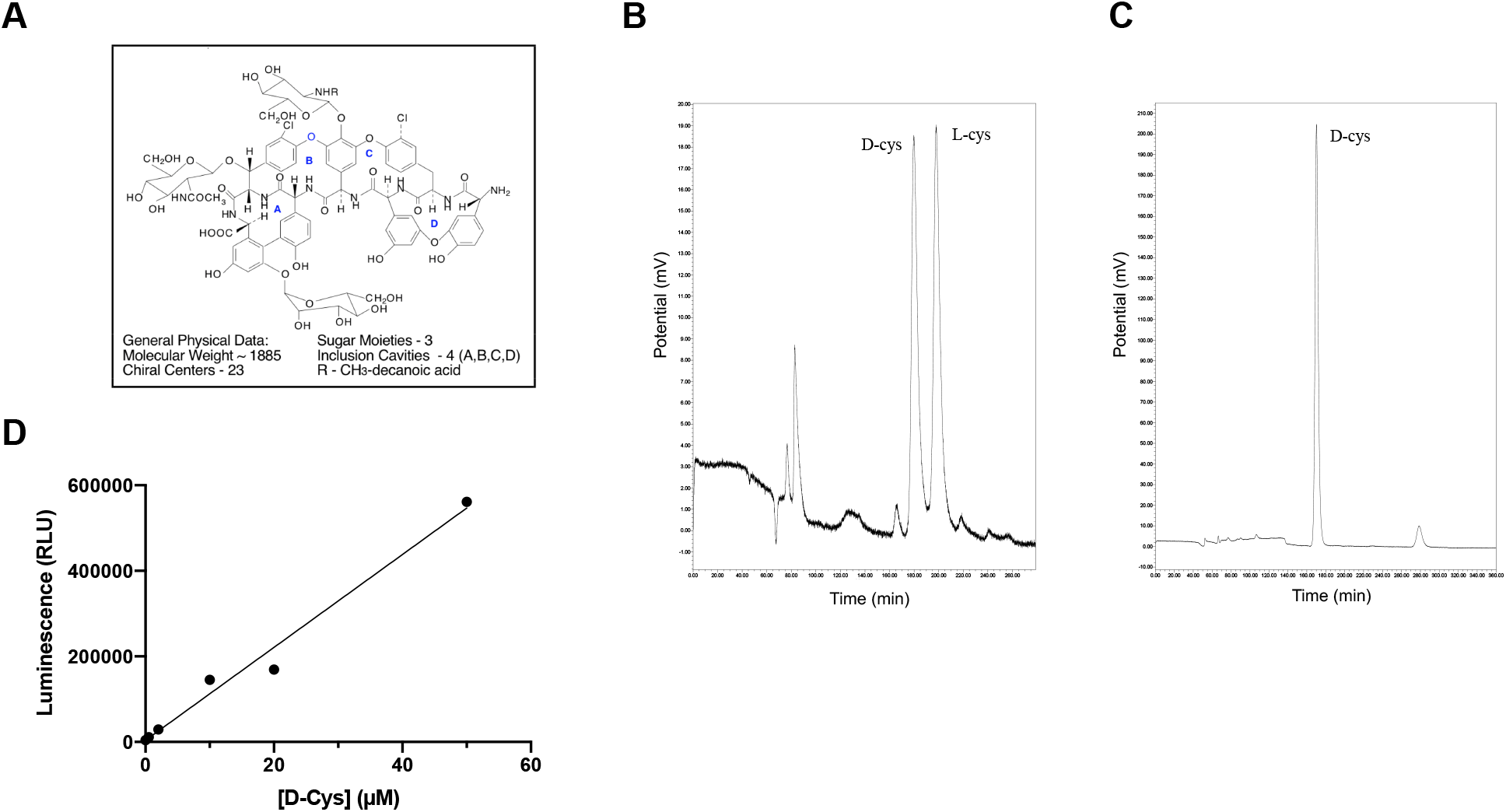
Measurement of free D-cysteine by high-performance liquid chromatography and luciferase assay. (*A*) Structure of the teicoplanin-based CHIROBIOTIC T chiral selector. (*B-C*) Representative chromatograms from HPLC analysis of (*B*) purified L-cys and D-cys standards (25 μM each) and (*C*) human cerebrospinal fluid. (*D*) D-cys luciferase assay standard curve. Known concentrations of D-cys (0.5-50 μM) were added to mouse brain lysates and analyzed by luciferase assay.

**Figure S2.**
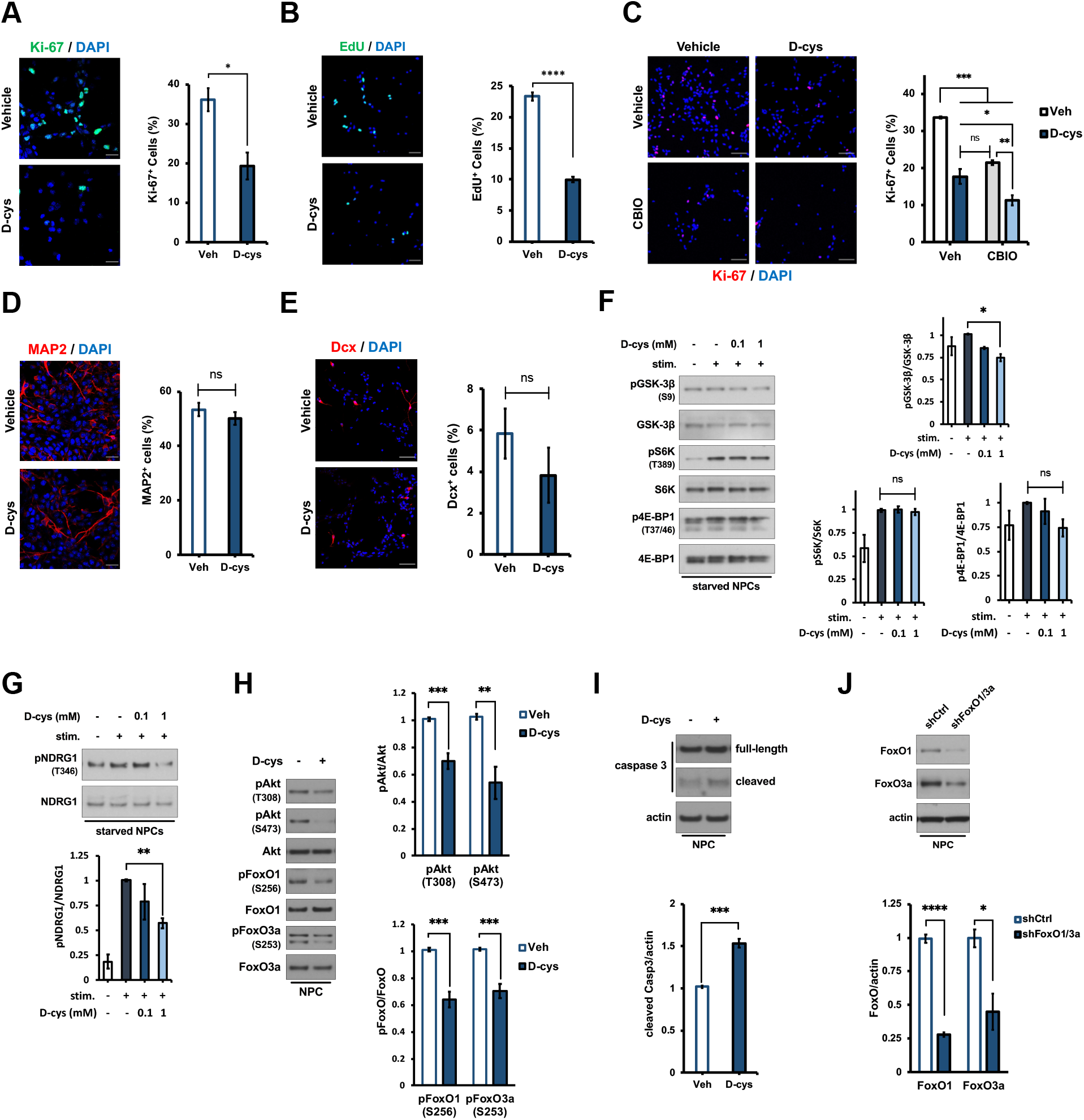
Antiproliferative actions of D-cysteine on NPCs involve Akt inhibition-dependent FoxO activation but not generation of hydrogen sulfide. (*A-B*) Representative images and quantification of (*A*) Ki-67, (*B*) EdU, and DAPI staining in NPCs treated with saline (vehicle) or 1 mM D-cys for 48 h (*n* = 3). Data were quantified as the number of Ki-67/EdU-positive (Ki-67^+^/EdU^+^) cells as a percent of the total number of (DAPI^+^) cells. Scale bar 20 μm (*A*) and 50 μm (*B*). (*C*) Immunostaining for Ki-67 and DAPI in NPCs treated with vehicle, 1 mM D-cys, and/or 5 μM CBIO for 48 h (*n* = 3). Data were quantified as in (*A*). Scale bar 50 μm. (*D-E*) Immunostaining for (*D*) MAP2, (*E*) doublecortin (Dcx), and DAPI in NPCs treated with vehicle or 1 mM D-cys for 48 h (*n* = 3). Data were quantified as in (*A*). Scale bar 20 μm (*D*) and 50 μm (*E*). (*F-G*) Representative immunoblots and quantification of (*F*) phospho- and total GSK-3β, S6K, and 4E-BP1 and (*G*) phospho- and total NDRG1 in NPCs. Cells were starved overnight followed by treatment with vehicle or the indicated concentration of D-cys for 1 h, followed by stimulation with complete medium for 5 minutes. Data are expressed as the normalized ratio of phosphorylated to total protein in each condition (*n* = 3). (*H-I*) Immunoblots and quantification of (*H*) phospho- and total Akt, FoxO1, and FoxO3a and (*I*) cleaved and total caspase 3 and actin in NPCs treated with vehicle or D-cys for 48 h (*n* = 3). (*J*) Immunoblots and quantification of FoxO1 and FoxO3a in NPCs infected with lentiviruses encoding non-targeting control (shCtrl) or FoxO1- and FoxO3a-targeting shRNAs (shFoxO1/3a) for 72 hours, normalized to actin (*n* = 3). Data are graphed as mean ± SEM. \**P* < 0.05, \*\**P* < 0.01, \*\*\**P* < 0.001, \*\*\*\**P* < 0.0001, ns = *P* > 0.05, two-tailed unpaired student’s *t*-test (*A-B, D-E, H-I*), or ANOVA with *post-hoc* Tukey’s test (*C, F, G*).

**Figure S3.**
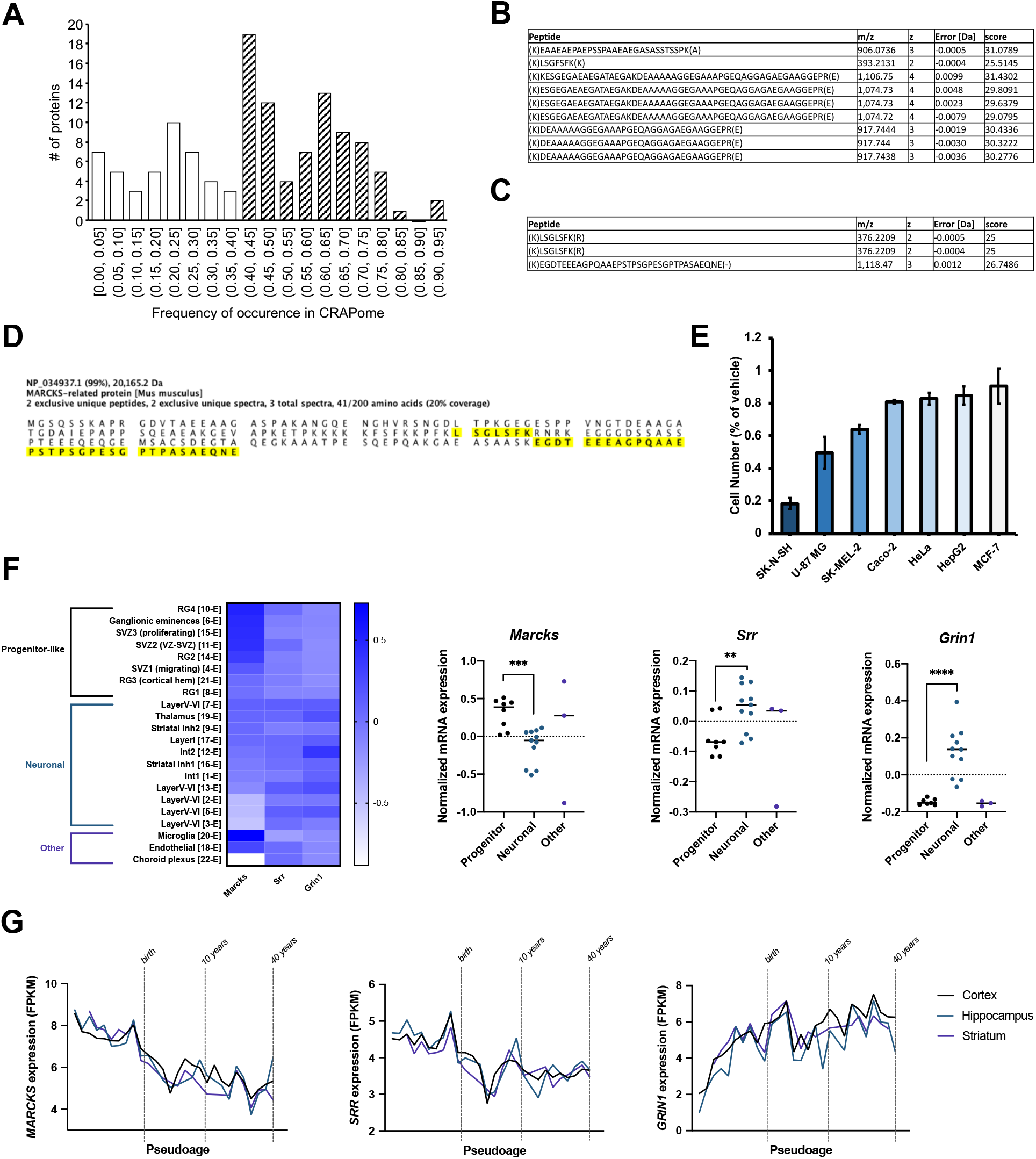
Mass spectrometric and bioinformatic analysis of MARCKS and MARCKSL1 as putative D-cysteine-binding proteins. (*A*) Histogram of proteins identified in mass spectrometric analysis of anti-D-cys immunoprecipitates but not preimmune IgG immunoprecipitates, sorted by frequency of occurrence in 716 affinity purification experiments registered in the CRAPome (**43**) (available at http://www.reprint-apms.org). Proteins with less than 40% occurrence in the CRAPome (white bars) were prioritized as likely true-positive hits. (*B-C*) Tables of peptides from (*B*) MARCKS and (*C*) MARCKSL1 in anti-D-cys immunoprecipitates. (*D*) Amino acid sequence of mouse MARCKSL1 protein. Peptides identified by mass spectrometric analysis of anti-D-cys immunoprecipitates are highlighted in yellow. (*E*) Antiproliferative effect of D-cys, as determined by the ratio of cell number after treatment with 1 mM D-cys for 48 h compared with vehicle-treated cells, in the indicated human cancer cell lines (*n* = 3). (*F*) Heatmap and graphs of single-cell RNA-sequencing data generated by Loo et. al (**45**) of *Marcks*, *Srr*, and *Grin1* levels in cell clusters of the E14.5 mouse brain, sorted into progenitor, neuronal, and other populations. (*G*) Analysis of *MARCKS*, *SRR*, and *GRIN1* mRNA in the developing human brain from the BrainSpan atlas (**46**) (available at https://www.brainspan.org).

## REFERENCES

1. J. J. Corrigan, D-amino acids in animals. Science 164, 142–149 (1969).

2. D. S. Dunlop, A. Neidle, D. McHale, D. M. Dunlop, A. Lajtha, The presence of free D-aspartic acid in rodents and man. Biochem. Biophys. Res. Commun. 141, 27–32 (1986).

3. S. H. Snyder, P. M. Kim, D-amino acids as putative neurotransmitters: Focus on D-serine. Neurochem. Res. 25, 553–560 (2000).

4. A. Hashimoto, et al., The presence of free D-serine in rat brain. FEBS Lett. 296, 33–36 (1992).

5. J.-P. Mothet, et al., D-Serine is an endogenous ligand for the glycine site of the N-methyl-D-aspartate receptor. Proc. Natl. Acad. Sci. U.S.A. 97, 4926–4931 (2000).

6. K. Erreger, et al., Subunit-specific agonist activity at NR2A-, NR2B-, NR2C-, and NR2D-containing N-methyl-D-aspartate glutamate receptors. Mol. Pharmacol. 72, 907–920 (2007).

7. M. J. Schell, O. B. Cooper, S. H. Snyder, D-aspartate localizations imply neuronal and neuroendocrine roles. Proc. Natl. Acad. Sci. U.S.A. 94, 2013–2018 (1997).

8. A. S. Huang, et al., D-aspartate regulates melanocortin formation and function: Behavioral alterations in D-aspartate oxidase-deficient mice. J. Neurosci. 26, 2814–2819 (2006).

9. H. Wolosker, S. Blackshaw, S. H. Snyder, Serine racemase: A glial enzyme synthesizing D-serine to regulate glutamate-N-methyl-D-aspartate neurotransmission. Proc. Natl. Acad. Sci. U.S.A. 96, 13409–13414 (1999).

10. P. M. Kim, et al., Aspartate racemase, generating neuronal D-aspartate, regulates adult neurogenesis. Proc. Natl. Acad. Sci. U.S.A. 107, 3175–3179 (2010).

11. H. A. Krebs, Metabolism of amino-acids: Deamination of amino-acids. Biochem. J. 29, 1620–1644 (1935).

12. L. Pollegioni, S. Sacchi, G. Murtas, Human D-amino acid oxidase: Structure, function, and regulation. Front. Mol. Biosci. 5 (2018).

13. T. Simonic, et al., cDNA cloning and expression of the flavoprotein D-aspartate oxidase from bovine kidney cortex. Biochem. J. 322, 729–735 (1997).

14. C. A. Weatherly, et al., D-amino acid levels in perfused mouse brain tissue and blood: A comparative study. ACS Chem. Neurosci. 8, 1251–1261 (2017).

15. E. H. Man, J. L. Bada, Dietary D-amino acids. Annu. Rev. Nutr. 7, 209–225 (1987).

16. I. Ilisz, A. Aranyi, Z. Pataj, A. Péter, Enantiomeric separation of nonproteinogenic amino acids by high-performance liquid chromatography. J. Chromatogr. A 1269, 94–121 (2012).

17. T. Toyo’oka, K. Imai, New fluorogenic reagent having halogenobenzofurazan structure for thiols: 4-(aminosulfonyl)-7-fluoro-2,1,3-benzoxadiazole. Anal. Chem. 56, 2461–2464 (1984).

18. H. H. Seliger, W. D. McElroy, E. H. White, G. F. Field, Stereospecificity and firefly bioluminescence, a comparison of natural and synthetic luciferins. Proc. Natl. Acad. Sci. U.S.A. 47, 1129–1134 (1961).

19. A. Kriegstein, A. Alvarez-Buylla, The glial nature of embryonic and adult neural stem cells. Annu. Rev. Neurosci. 32, 149–184 (2009).

20. N. D. Dwyer, et al., Neural stem cells to cerebral cortex: Emerging mechanisms regulating progenitor behavior and productivity. J. Neurosci. 36, 11394–11401 (2016).

21. D. R. Kaplan, K. Matsumoto, E. Lucarelli, C. J. Thielet, Induction of TrkB by retinoic acid mediates biologic responsiveness to BDNF and differentiation of human neuroblastoma cells. Neuron 11, 321–331 (1993).

22. B. Laurent, et al., A specific LSD1/KDM1A isoform regulates neuronal differentiation through H3K9 demethylation. Mol. Cell 57, 957–970 (2015).

23. N. Shibuya, et al., A novel pathway for the production of hydrogen sulfide from D-cysteine in mammalian cells. Nat. Commun. 4 (2013).

24. D. Ferraris, et al., Synthesis and biological evaluation of D-amino acid oxidase inhibitors. J. Med. Chem. 51, 3357–3359 (2008).

25. B. D. Manning, L. C. Cantley, AKT/PKB signaling: Navigating downstream. Cell 129, 1261–1274 (2007).

26. D. R. Alessi, et al., Mechanism of activation of protein kinase B by insulin and IGF-1. EMBO J. 15, 6541–6551 (1996).

27. A. Brunet, et al., Akt promotes cell survival by phosphorylating and inhibiting a forkhead transcription factor. Cell 96, 857–868 (1999).

28. D. A. E. Cross, D. R. Alessi, P. Cohen, M. Andjelkovich, B. A. Hemmings, Inhibition of glycogen synthase kinase-3 by insulin mediated by protein kinase B. Nature 378, 785–789 (1995).

29. Y. Sancak, et al., PRAS40 is an insulin-regulated inhibitor of the mTORC1 protein kinase. Mol. Cell 25, 903–915 (2007).

30. P. E. Burnett, R. K. Barrow, N. A. Cohen, S. H. Snyder, D. M. Sabatini, RAFT1 phosphorylation of the translational regulators p70 S6 kinase and 4E-BP1. Proc. Natl. Acad. Sci. U.S.A. 95, 1432–1437 (1998).

31. A. Brunet, et al., Protein kinase SGK mediates survival signals by phosphorylating the forkhead transcription factor FKHRL1 (FOXO3a). Mol. Cell. Biol. 21, 952–965 (2001).

32. J. T. Murray, et al., Exploitation of KESTREL to identify NDRG family members as physiological substrates for SGK1 and GSK3. Biochem. J. 384, 477–488 (2004).

33. B. M. T. Burgering, P. J. Coffer, Protein kinase B (c-Akt) in phosphatidylinositol-3-OH kinase signal transduction. Nature 376, 599–602 (1995).

34. R. H. Medema, G. J. P. L. Kops, J. L. Bos, B. M. T. Burgering, AFX-like Forkhead transcription factors mediate cell-cycle regulation by Ras and PKB through p27kip1. Nature 404, 782–787 (2000).

35. X. Zhang, et al., FOXO1 is an essential regulator of pluripotency in human embryonic stem cells. Nat. Cell Biol. 13, 1092–1099 (2011).

36. V. Graham, J. Khudyakov, P. Ellis, L. Pevny, SOX2 functions to maintain neural progenitor identity. Neuron 39, 749–765 (2003).

37. A. J. Blaschke, K. Staley, J. Chun, Widespread programmed cell death in proliferative and postmitotic regions of the fetal cerebral cortex. Development 122, 1165–1174 (1996).

38. K. Kuida, et al., Decreased apoptosis in the brain and premature lethality in CPP32-deficient mice. Nature 384, 368–372 (1996).

39. K. Kuida, et al., Reduced apoptosis and cytochrome c–mediated caspase activation in mice lacking caspase 9. Cell 94, 325–337 (1998).

40. R. F. Hevner, et al., Tbr1 regulates differentiation of the preplate and layer 6. Neuron 29, 353–366 (2001).

41. P. Arlotta, et al., Neuronal subtype-specific genes that control corticospinal motor neuron development in vivo. Neuron 45, 207–221 (2005).

42. O. Britanova, S. Akopov, S. Lukyanov, P. Gruss, V. Tarabykin, Novel transcription factor Satb2 interacts with matrix attachment region DNA elements in a tissue-specific manner and demonstrates cell-type-dependent expression in the developing mouse CNS. Eur J. Neurosci. 21, 658–668 (2005).

43. D. Mellacheruvu, et al., The CRAPome: a contaminant repository for affinity purification–mass spectrometry data. Nat. Methods 10, 730–736 (2013).

44. M. Glaser, et al., Myristoylated alanine-rich C kinase substrate (MARCKS) produces reversible inhibition of phospholipase C by sequestering phosphatidylinositol 4,5-bisphosphate in lateral domains. J. Biol. Chem. 271, 26187–26193 (1996).

45. L. Loo, et al., Single-cell transcriptomic analysis of mouse neocortical development. Nat. Commun. 10 (2019).

46. J. A. Miller, et al., Transcriptional landscape of the prenatal human brain. Nature 508, 199–206 (2014).

47. D. J. Stumpo, J. M. Graff, K. A. Albert, P. Greengard, P. J. Blackshear, Molecular cloning, characterization, and expression of a cDNA encoding the “80- to 87-kDa” myristoylated alanine-rich C kinase substrate: a major cellular substrate for protein kinase C. Proc. Natl. Acad. Sci. U.S.A. 86, 4012–4016 (1989).

48. M. Thelen, A. Rosen, A. C. Nairn, A. Aderem, Regulation by phosphorylation of reversible association of a myristoylated protein kinase C substrate with the plasma membrane. Nature 351, 320–322 (1991).

49. B. P. Ziemba, J. E. Burke, G. Masson, R. L. Williams, J. J. Falke, Regulation of PI3K by PKC and MARCKS: Single-molecule analysis of a reconstituted signaling pathway. Biophys. J. 110, 1811–1825 (2016).

50. M. J. Schell, M. E. Molliver, S. H. Snyder, D-serine, an endogenous synaptic modulator: localization to astrocytes and glutamate-stimulated release. Proc. Natl. Acad. Sci. U.S.A. 92, 3948–3952 (1995).

51. L.-Z. Wang, X.-Z. Zhu, Spatiotemporal relationships among D-serine, serine racemase, and D-amino acid oxidase during mouse postnatal development. Acta Pharmacol. Sin. 24, 965–974 (2003).

52. A. Hashimoto, et al., Embryonic development and postnatal changes in free D-aspartate and D-serine in the human prefrontal cortex. J. Neurochem. 61, 348–351 (1993).

53. R. Raballo, et al., Basic fibroblast growth factor (Fgf2) is necessary for cell proliferation and neurogenesis in the developing cerebral cortex. J. Neurosci. 20, 5012–5023 (2000).

54. M. Groszer, et al., Negative regulation of neural stem/progenitor cell proliferation by the Pten tumor suppressor gene in vivo. Science 294, 2186–2189 (2001).

55. J. D. Buxbaum, et al., Mutation screening of the PTEN gene in patients with autism spectrum disorders and macrocephaly. Am. J. Med. Genet. 144B, 484–491 (2007).

56. A. M. D’Gama, et al., Somatic mutations activating the mTOR pathway in dorsal telencephalic progenitors cause a continuum of cortical dysplasias. Cell Rep. 21, 3754–3766 (2017).

57. V. Stambolic, et al., Negative Regulation of PKB/Akt-Dependent Cell Survival by the Tumor Suppressor PTEN. Cell 95, 29–39 (1998).

58. D. Jeong, et al., LRIG1-mediated inhibition of EGF receptor signaling regulates neural precursor cell proliferation in the neocortex. Cell Rep. 33, 108257 (2020).

59. J. Paik, et al., FoxOs cooperatively regulate diverse pathways governing neural stem cell homeostasis. Cell Stem Cell 5, 540–553 (2009).

60. V. M. Renault, et al., FoxO3 regulates neural stem cell homeostasis. Cell Stem Cell 5, 527–539 (2009).

61. D. J. Stumpo, C. B. Bock, J. S. Tuttle, P. J. Blackshear, MARCKS deficiency in mice leads to abnormal brain development and perinatal death. Proc. Natl. Acad. Sci. U.S.A. 92, 944–948 (1995).

62. J. M. Weimer, et al., MARCKS modulates radial progenitor placement, proliferation and organization in the developing cerebral cortex. Development 136, 2965–2975 (2009).

63. A. K. Mustafa, et al., Glutamatergic regulation of serine racemase via reversal of PIP2 inhibition. Proc. Natl. Acad. Sci. U.S.A. 106, 2921–2926 (2009).

64. D. J. Laurie, P. H. Seeburg, Regional and developmental heterogeneity in splicing of the rat brain NMDAR1 mRNA. J. Neurosci. 14, 3180–3194 (1994).

65. J. B. Monihan, V. M. Corpus, W. F. Hood, J. W. Thomas, R. P. Compton, Characterization of a [3H]glycine recognition site as a modulatory site of the N-methyl-D-aspartate receptor complex. J. Neurochem. 53, 370–375 (1989).

66. K. Abe, H. Kimura, The possible role of hydrogen sulfide as an endogenous neuromodulator. J. Neurosci 16, 1066–1071 (1996).

67. T. Seki, et al., D-cysteine promotes dendritic development in primary cultured cerebellar Purkinje cells via hydrogen sulfide production. Mol. Cell. Neurosci. 93, 36–47 (2018).

68. A. Y. Goltsov, et al., Polymorphism in the 5′-promoter region of serine racemase gene in schizophrenia. Mol. Psychiatry 11, 325–326 (2006).

69. Y. Morita, et al., A genetic variant of the serine racemase gene is associated with schizophrenia. Biol. Psychiatry 61, 1200–1203 (2007).

70. V. Labrie, et al., Serine racemase is associated with schizophrenia susceptibility in humans and in a mouse model. Hum. Mol. Genet. 18, 3227–3243 (2009).

71. I. Bendikov, et al., A CSF and postmortem brain study of D-serine metabolic parameters in schizophrenia. Schizophr. Res. 90, 41–51 (2007).

72. D. T. Balu, et al., Multiple risk pathways for schizophrenia converge in serine racemase knockout mice, a mouse model of NMDA receptor hypofunction. Proc. Natl. Acad. Sci. U.S.A. 110, E2400–E2409 (2013).

73. D. T. Balu, J. T. Coyle, The NMDA receptor ‘glycine modulatory site’ in schizophrenia: D-serine, glycine, and beyond. Curr. Opin. Pharmacol. 20, 109–115 (2015).

74. S. Gulsuner, et al., Spatial and temporal mapping of de novo mutations in schizophrenia to a fetal prefrontal cortical network. Cell 154, 518–529 (2013).

75. A. E. Jaffe, et al., Developmental and genetic regulation of the human cortex transcriptome illuminate schizophrenia pathogenesis. Nat. Neurosci. 21, 1117–1125 (2018).

76. L. D. Selemon, N. Zecevic, Schizophrenia: a tale of two critical periods for prefrontal cortical development. Transl. Psychiatry 5, e623 (2015).

77. R. Birnbaum, D. R. Weinberger, Genetic insights into the neurodevelopmental origins of schizophrenia. Nat. Rev. Neurosci. 18, 727–740 (2017).

78. A. L. Pinner, V. Haroutunian, J. H. Meador-Woodruff, Alterations of the myristoylated, alanine-rich C kinase substrate (MARCKS) in prefrontal cortex in schizophrenia. Schizophr. Res. 154, 36–41 (2014).

79. L. M. Huckins, et al., Gene expression imputation across multiple brain regions provides insights into schizophrenia risk. Nat. Genet. 51, 659–674 (2019).

80. P. J. Thul, et al., A subcellular map of the human proteome. Science 356 (2017).

81. A. C. Basu, et al., Targeted disruption of serine racemase affects glutamatergic neurotransmission and behavior. Mol. Psychiatry 14, 719–727 (2009).

82. M. M. Harraz, et al., Cocaine-induced locomotor stimulation involves autophagic degradation of the dopamine transporter. Mol Psychiatry 26, 370–382 (2021).

83. E. Abramson, et al., Designed PKC-targeting bryostatin analogs modulate innate immunity and neuroinflammation. Cell Chem. Biol. 28, 537–545 (2021).

84. M. M. Harraz, et al., Cocaine receptor identified as BASP1. bioRxiv doi:10.1101/2020.11.23.392787 (2020).

85. P. Guha, et al., Loss of PI3-kinase activity of inositol polyphosphate multikinase impairs PDK1-mediated AKT activation, cell migration, and intestinal homeostasis. bioRxiv doi:10.1101/2020.12.18.423145 (2020).

